# Uncovering critical transitions and molecule mechanisms in disease progressions using Gaussian graphical optimal transport

**DOI:** 10.1101/2024.04.24.590914

**Authors:** Wenbo Hua, Ruixia Cui, Heran Yang, Jingyao Zhang, Chang Liu, Jian Sun

**Affiliations:** School of Mathematics and Statistics, Xi’an Jiaotong University, No.28 Xianning West Rd., Xi’an, 710049, Shaanxi, China; Key Laboratory of Surgical Critical Care and Life Support, Xi’an Jiaotong University Ministry of Education, No.28 Xianning West Rd., Xi’an, 710049, Shaanxi, China; Department of Hepatobiliary Surgery and Liver Transplantation, The Second Affiliated Hospital of Xi’an Jiaotong University, No.154 West 5th Rd., Xi’an, 710004, Shaanxi, China; Department of SICU, The First Affiliated Hospital of Xi’an Jiaotong University, No.227 Yanta West Rd., Xi’an, 710061, Shaanxi, China

**Keywords:** Disease Progression Analysis, Gaussian Graphical Model, Optimal Transport, Critical Transitions Detection, Trigger Molecules Identification, Single Sample Prediction

## Abstract

Understanding disease progression is crucial for detecting critical transitions and finding trigger molecules, facilitating early diagnosis interventions. However, the high dimensionality of data and the lack of aligned samples across disease stages have posed challenges in addressing these tasks. We present a novel framework, Gaussian Graphical Optimal Transport (GGOT), for analyzing disease progressions. The proposed GGOT uses Gaussian graphical models, incorporating protein interaction networks, to characterize the data distributions at different disease stages. Then we use population-level optimal transport to calculate the Wasserstein distances and transport maps between stages, enabling us to detect critical transitions. By analyzing the per-molecule transport distance, we quantify the importance of each molecule and identify trigger molecules. Moreover, GGOT predicts the occurrence of critical transitions in unseen samples and visualizes the disease progression process. We apply GGOT to the simulation dataset and six disease datasets with varying disease progression rates, to show its effectiveness for detecting critical transitions and identifying trigger molecules.

## 1 Introduction

It is known that disease progression is a dynamic process and may consist of several pathological stages [1]. The transitions between different stages reflect the disease evolution and the underlying triggering factors, which help disease treatment and healthcare [2]. Numerous evidences indicate that despite the high variability in the progression of different diseases, many diseases can exhibit similar sudden deterioration across progression, such as epileptic seizures [3, 4], asthma attacks [5] and prostate cancer [6]. The phenomenon of the sudden deterioration in disease is called critical transition, and the corresponding threshold when the critical transition occurs is called the tipping point [7], which exists in many complex systems [8]. The systems can suddenly shift from one state to another at tipping points [9]. Through the critical transition, the progression of different diseases can be categorized into three states, i.e., normal, critical, and disease state (Fig. 1a). The critical state serves as the pre-transition state near the tipping point before disease appearance [10–12]. That is, when approaching the tipping point, the patient typically undergoes a disastrous and irreversible transition, usually manifesting as deterioration [3–5]. Before reaching the critical state, disease progression is slow, and there is no discernible difference in its clinical presentation compared to the normal stage. The appropriate treatment can restore the system to its normal state. But after reaching the tipping point, the state can not be regained [12]. The irreversible deterioration of diseases can seriously threaten the life and health of patients. Therefore, comprehending disease progressions and detecting critical transitions are important for assessing the state stability of patients, assisting in early diagnosis and treatment [9, 13].

**Fig. 1:**
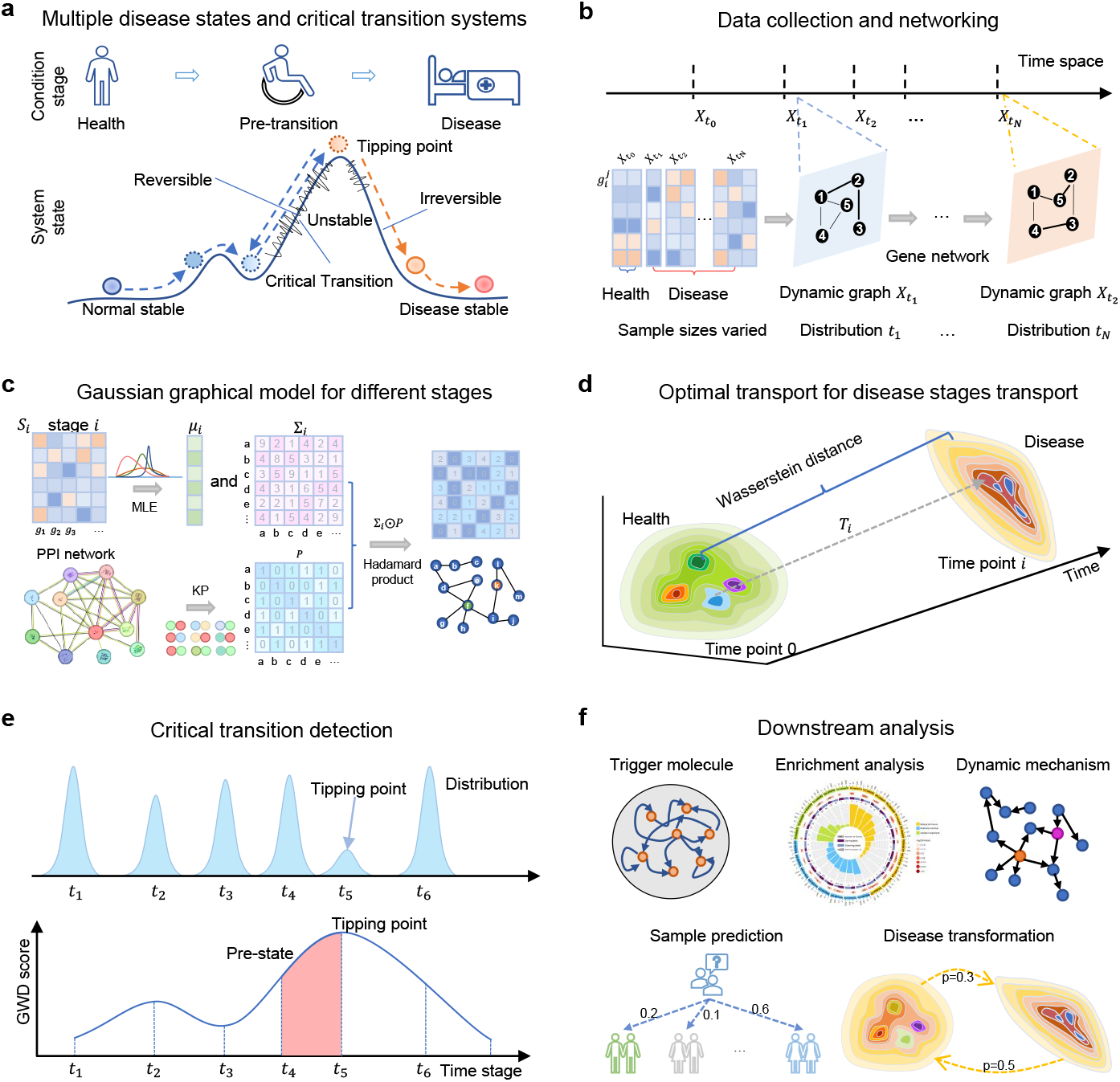
Overview of the GGOT model. **a** The critical transition phenomena always exhibit in disease progressions. Disease progression stages can be categorized into three states, i.e., normal, critical, and disease state, where the critical state indicates the potential for irreversible deterioration of patients. **b** According to the pathological characteristics, the disease samples from different stages are collected for graph modeling. The sample sizes vary typically for each stage. **c** The GGOT model utilizes prior knowledge from PPI networks and maximum likelihood estimation to establish Gaussian graphical distributions, characterizing the disease state for each stage. Distributions reflect dependencies between genes. **d** For a abnormal stage *i*, we aim to model it with a function *T*_*i*_ that maps distribution from normal stage 0 to abnormal stage *i*. In particular, we calculate *T*_*i*_ through the proposed Gaussian optimal transport due to a lack of paired measurements. **e** Detecting tipping points by Wasserstein distance between normal stage 0 and abnormal stage *i*. The critical transition appears at the stage corresponding to the maximum Wasserstein distance. **f** Downstream analysis. We perform downstream validation analysis of the results, including identifying trigger molecules, functional analysis, sample prediction, and so on.

However, it is challenging to address this issue. The state of patients remains stable and may show little change before reaching the tipping point, resulting in such transitions being abrupt and difficult to recognize [10–12, 14]. In addition, system noise, patient heterogeneity, sample imbalances, the lack of aligned samples, and model inaccuracy hinder reliable detection of the critical transition. Patients with the same disease diagnosis may exhibit varying early symptoms and developmental trajectories [15, 16], responses diversely to identical risk factors and treatments [17]. Indeed, disease development and progression are derived by the concerted actions of numerous molecules, with a highly complex genetic pattern [18]. Gene networks will exhibit on-off nonlinear behavior in response to internal signals or external stimuli [19]. Among all molecules, there are always certain key genes or proteins that play a crucial role in disease progression, e.g., in determining critical transitions. These key molecules can serve as biomarkers for disease diagnosis, prediction, and treatment. Identifying trigger molecules while detecting critical transitions offers greater insight into the mechanisms of disease progression. With the development of high-throughput sequencing technology, it has become simple to obtain gene expression data [20]. Gene expression data provides an excellent and abundant source of information for investigating dynamic activities [21]. Gene regulatory networks determined by gene expression data are often used to represent the dynamic networks of diseases to study disease progression [22, 23]. Previous studies have demonstrated that critical transitions in disease progression may have some generic properties regardless of differences between diseases [11, 24]. The key to solving this issue fundamentally hinges on seeking these generic properties.

Recent studies to detect critical transitions of diseases can be mainly categorized into two categories. The first category is based on the flux theory upon the nonequilibrium dynamics [25]. This theory calculates the probability of the system remaining at each point in the state space by reconstructing the system quasi-potential and deducing various properties of the system state transitions [26]. Although the theoretical basis of this method is reliable, its computational complexity is high, making it difficult to apply to large-scale, high-dimensional biological systems [27]. The second category is based on the theory of “critical slowing down” using the elasticity model of dynamic systems [19, 28]. Critical slowing down refers to the phenomenon in which a system gradually slows down while recovering from a slightly disrupted state to approach equilibrium near the tipping point [9]. This theory is widely utilized, but it requires extended periods of high-resolution data for estimation [29] and may not be reliable in identifying tipping points [30]. The dynamic network biomarkers (DNB) developed by critical slowing down [11, 12, 24, 31–33] explore a model-free statistical approach, lessening constraints on data sampling. It can not be employed for noisy systems or those with fast state transitions [34], and does not provide more information about progressions and molecular mechanisms. In brief, previous methods are limited to addressing specific problems in complex diseases and can not tackle those that are highly unbalanced, noisy, and exhibit rapid transition rates. Our objective is to present a universal framework for detecting critical transitions, that can address these limitations.

In this work, we propose the *Gaussian Graphical Optimal Transport* (GGOT), a novel and generalized approach that understands disease progression and detects critical transitions by directly learning maps and uncovering the relative distance from normal to disease stages, further clarifying the molecular changes and mechanism of progression of the critical transition of diseases. Assuming the progression of diseases gradually alters the gene interaction networks of patients, we characterize the interactions between genes in terms of Gaussian graphical model embedded with prior knowledge of biomolecular networks (Protein-Protein Interaction Network, PPI [35]), and learn the distribution changes using optimal transport (OT) theory [36]. The Gaussian graphical model is a probabilistic graphical model for describing dependencies between random variables [37], and is frequently used to analyze gene expression data, aiding in the uncovering of interactions and regulatory relationships between genes [38, 39]. The changes among distinct Gaussian graphical distributions can be modeled through the principles of optimal transport. Optimal transport is a theory that describes the optimal way to transfer data between two different distributions [36] and has recently been applied to genomics data analysis. It has shown great superiority in problems such as spatial annotation of scRNA-seq data [40], transcriptome data alignment and integration [41], and cell-cell communication [42]. However, previous OT-based methods can not detect critical transitions of diseases and provide interpretability under actual molecule interaction relationships.

Based on developments of Gaussian graphical model and optimal transport, GGOT constructs different disease stages using Gaussian graphical model embedded with PPI, to reduce the influence of irrelevant variables and enhance the interpretability of the model by considering the actual biomolecular association. Then, GGOT learns the optimal transport map for each disease stage and uses the Wasserstein distance to measure the relative distance between different disease stages. This distance can serve as an early warning signal for tipping point prediction, because it quantifies the minimum “effort” needed to transition between disease stages, sensitive to gene component changes. Besides, GGOT can identify trigger molecules by decomposing the Wasserstein distance, determine biomarkers, reveal the key pathways and functions, predict the distribution of disease states in unknown samples, and visualize the disease transport process, which are significant to early healthcare.

We demonstrate the effectiveness of GGOT using simulation data and six complex real-world disease datasets with varying progression rates. The results show that GGOT can robustly handle real disease data, precisely detect tipping points across diverse disease categories, and identify trigger molecules at tipping points. Moreover, it promotes the discovery of biomarkers and regulatory mechanisms between different trigger molecules and reveals trajectories of disease progression. Finally, we collect samples from patients with sepsis to investigate the phenomenon of critical transitions in acute progressive disease. Our study yields significant results by identifying new trigger molecules, emphasizing that GGOT remains effective in acute progressive diseases. These findings further illustrate the superiority of our approach.

## 2 Results

### 2.1 Overview of GGOT

Given temporal gene expression data 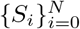 about the specific disease with *N*+1 disease stages, where 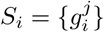 denotes the set of disease samples of stage *i*. We assume Ω_*S*_ = {0, 1, …, *N*} denotes all stages of the disease, with *i* = 0 as the normal stage and *i* ≠ 0 as the following abnormal stages. Critical transitions in biological systems are always accompanied by sudden changes in temporal signals, such as the significant increase in variance and auto-correlation coefficient [9]. The key to detecting critical transitions is defining a function that can quantitatively measure the changes in the system state with disease progression. Dynamic progressions of diseases are characterized by multi-criticality, instability, and high complexity (Fig. 1a). Therefore, this function must encapsulate the common underlying mechanisms of critical transitions across various diseases. In our approach, we introduce GGOT to quantify the relative distance from a healthy state to different disease states as the measure, which solves the challenges caused by sample imbalance and the lack of aligned samples and is robust to patient heterogeneity and system noise.

The proposed model GGOT has three features. First, we use the Gaussian graphical model (Fig. 1b) embedded with domain prior knowledge of PPI networks (Fig. 1c) to describe gene interaction networks and model data distributions in different disease stages, relieving the sample imbalance issue. Second, we model the corresponding optimal transport processes from normal to abnormal stages by the direction of disease progressions (Fig. 1d) and calculate the corresponding global Wasserstein distance to represent the minimum “effort” required to transition from normal to abnormal stages (Fig. 1e), to detect critical transitions of diseases. Last, we propose the local Wasserstein distance based on optimal transport decomposition to identify trigger molecules during critical transitions, determine biomarkers, and infer the mechanisms of disease exacerbation. We further develop downstream analytical tools to investigate the interaction relationships among molecules, predict whether an unknown sample reaches tipping points, and describe the transport processes at different stages of the disease (Fig. 1f).

According to Gaussian graphical optimal transport, we can detect critical transitions in diseases by solving a maximization problem, i.e.,

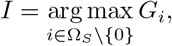

where 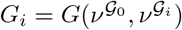, *i* = 1, 2, …, *N* is the **Global Wasserstein Distance** (GWD) score from stage 0 to *i* determined by Eq. (9). *T*_*i*_ is the corresponding optimal transport map, describing the transition process. The GWD score is sensitive to compositional differences that induce global changes in the graph structure. We further decompose the GWD score *G*_*i*_ into the **Local Wasserstein Distance** (LWD) score *L*_*i*_ of each specific gene by Eq. (11), where *L*_*i*_(*j*) measures the interaction difference of gene *j* under global effects. The difference in LWD scores reflects not only the changes in interactions associated with gene *j* but also the global structural significance of gene *j* (Supplementary Fig. 1). This characteristic helps to identify regulatory genes in key pathways, which is not available in previous methods.

To assess the effectiveness of the proposed method GGOT, we use a simulation dataset and six complex real-world disease datasets. The simulation dataset is generated using gene regulatory networks to simulate disease progression accurately. The complex disease datasets (Supplementary Fig. 2) are from GEO database and TCGA database, encompassing diseases with varying rates of progression and levels of severity (Supplementary Table 3). Dimensionality reduction methods can not capture the state transition properties (Supplementary Fig. 3). Details of the datasets, such as description, pathologic stages, and sample size, are provided in Supplementary Section C. For each dataset, GGOT detects disease critical transitions first and quantifies the transport distances from normal to abnormal stages (Section 2.3). Secondly, GGOT identifies trigger molecules at tipping points and performs functional analysis and survival analysis of identified molecules to illustrate regulatory mechanisms and biomarkers (Section 2.4). GGOT then predicts the stage distributions of the unknown sample, and determines whether the sample reaches the tipping point (Section 2.5). Finally, GGOT models the transport processes from normal to abnormal stages of diseases by optimal transport mapping and dimensionality reduction methods, such as PCA [43] and T-SNE [44]. The results visualize the global transport processes of stages distributions and the molecule transport processes in disease progressions (Section 2.6).

### 2.2 GGOT for simulation data validation

To validate the proposed method GGOT, we employ a gene regulatory network in the form of Michaelis-Menten or Hill with sixteen nodes to simulate the disease progression [24]. Regulatory networks employing Michaelis-Menten or Hill bifurcation are frequently utilized to depict the gene regulatory activities in biological systems, such as transcription, translation [45, 46]. These networks are instrumental in detecting critical transitions. The equation for the regulatory network (Fig. 2a) employed in our study is detailed in Supplementary Section D. The critical signal of the network is controlled by parameter *q*, with *q* = 0 as the bifurcation marking the tipping point. Furthermore, Nodes 1 to 7 are directly regulated by parameter *q* and are trigger molecules, while the remaining nodes are irrelevant molecules, independent of parameter *q*. Based on the variation of the parameter *q* from − 0.3 to 0.2, the dataset of numerical simulation is generated to show the effectiveness of detection using GGOT when the system approaches the tipping point.

**Fig. 2:**
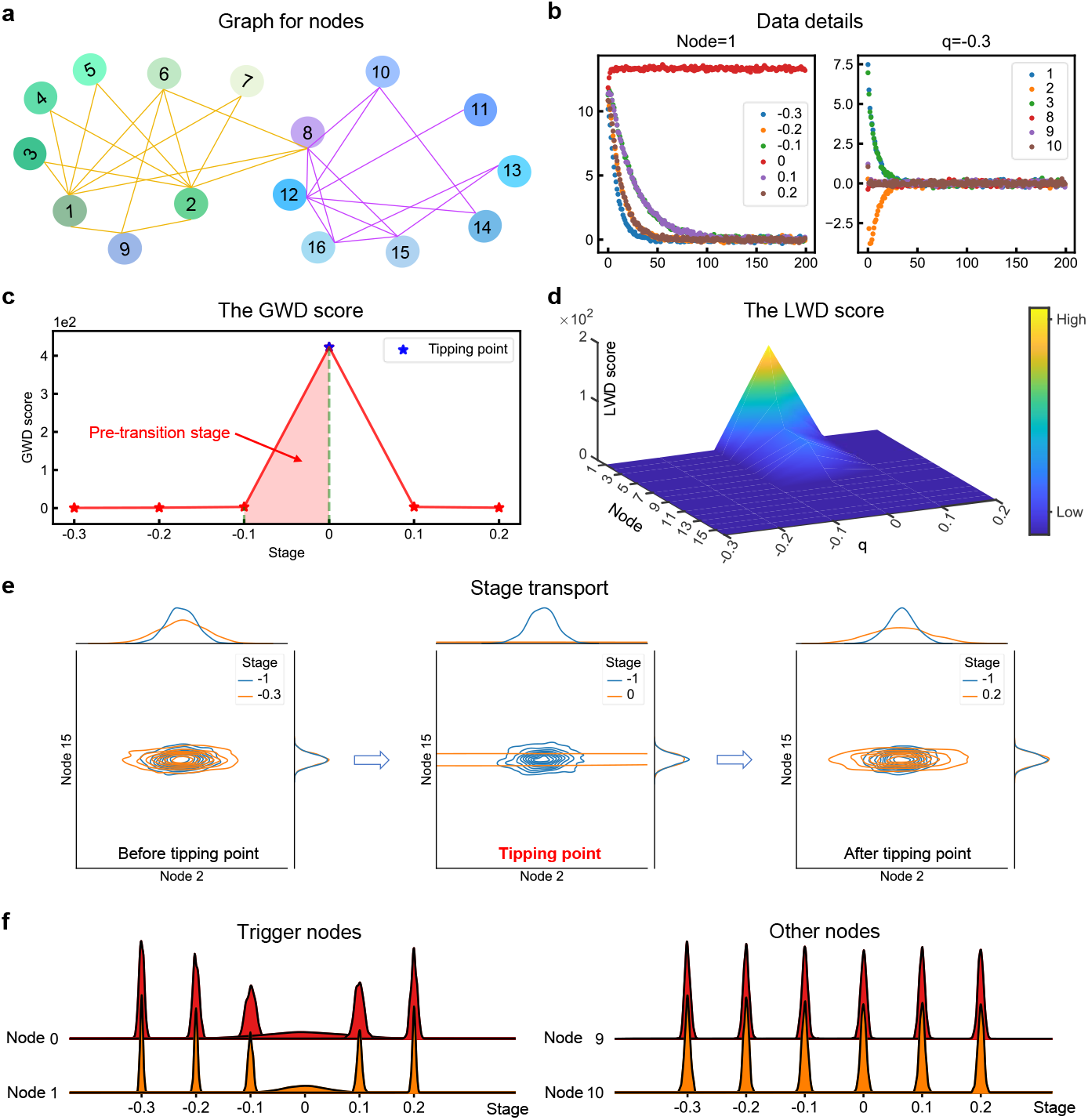
GGOT for simulation data validation. **a** The numerical simulation is conducted based on the graph with 16 nodes. The graph designed by the gene regulatory network reflects the relationships between nodes. **b** Data summary of nodes across varying conditions regulated by parameter *q*. The parameter *q* is the governing factor for the regulatory network in Supplementary Equation (S1). The left is the evolution of node 1 for different parameters *q*, and the right is the evolution of different nodes for *q* = − 0.3. **c** The curve of GWD score. The sudden increase in GWD scores heralds the coming of critical transition *q* = 0. **d** The landscape of LWD score. LWD scores reflect changes in single nodes within the system. The LWD scores of certain nodes, termed trigger molecules, exhibit a sudden increase as they approach the tipping point. **e** The changes of joint Gaussian distribution in nodes 2 and 15 between the normal stage (*q* = − 1) and other stages (− 0.3 ≤ *q* 0 ≤.2). Optimal transport can accurately calculate the changes in distributions of molecular synergy. **f** The comparison of the distribution progression between trigger molecules and other molecules at different *q*. Trigger molecules are more unstable than other molecules when *q* = 0.

In Fig. 2b, we demonstrate the changes in the value of node 1 under different parameter *q* settings and the variations in different nodes under the parameter *q* = − 0.3. It shows that parameter *q* controls the evolution of node values. The GWD score is displayed in Fig. 2c. It is seen that a sudden increase of GWD scores near the bifurcation *q* = 0 indicates the upcoming tipping point. When away from the tipping point, the GWD scores maintain a low level. To exhibit the distinct dynamics of each node in the system and identify trigger molecules during progression, we present the dynamics landscape of LWD scores in Fig. 2d. When the system is distant from the tipping point, all LWD scores are smooth and at a low level. As the system approaches the tipping point *q* = 0, the LWD scores of certain nodes (1, 2, …, 7) increase drastically, called trigger molecules, while the others remain low. It proves that GGOT can identify trigger molecules, and corresponding results are consistent with the nodes regulated by *q*. The distribution transport processes from normal to abnormal stages are illustrated in Fig. 2e. The system distribution gradually diverges and fluctuates more as it approaches the tipping point, and then returns to normal level as it moves away from the tipping point. We show the changes in marginal distributions of trigger molecules and other molecules in Fig. 2f. It indicates that the variance of the trigger molecules increases dramatically approaching the tipping point. Through the simulation experiment, it is demonstrated that GGOT is reliable and accurate in detecting critical transitions. Besides, GGOT can identify trigger molecules and explain critical changes in the progression of systems.

### 2.3 GGOT detects critical transitions in disease progression, uncovering the occurrence of irreversible changes

We apply GGOT to detect critical transitions for real-world disease datasets with varying progression rates (Supplementary Table 3). GGOT can identify patterns of the alterations in gene relationships during disease progression, allowing for early detection of tipping points for disease state transitions. We evaluate the effectiveness of GGOT for detecting tipping points in chronic and acute progressive diseases. The results of GSE48452, GSE2565, LUAD, and XJTUSepsis are shown in Figs. 3a-d, and the results of COAD, GSE154918 are shown in Supplementary Figs. 4a and 5a, respectively.

**Fig. 3:**
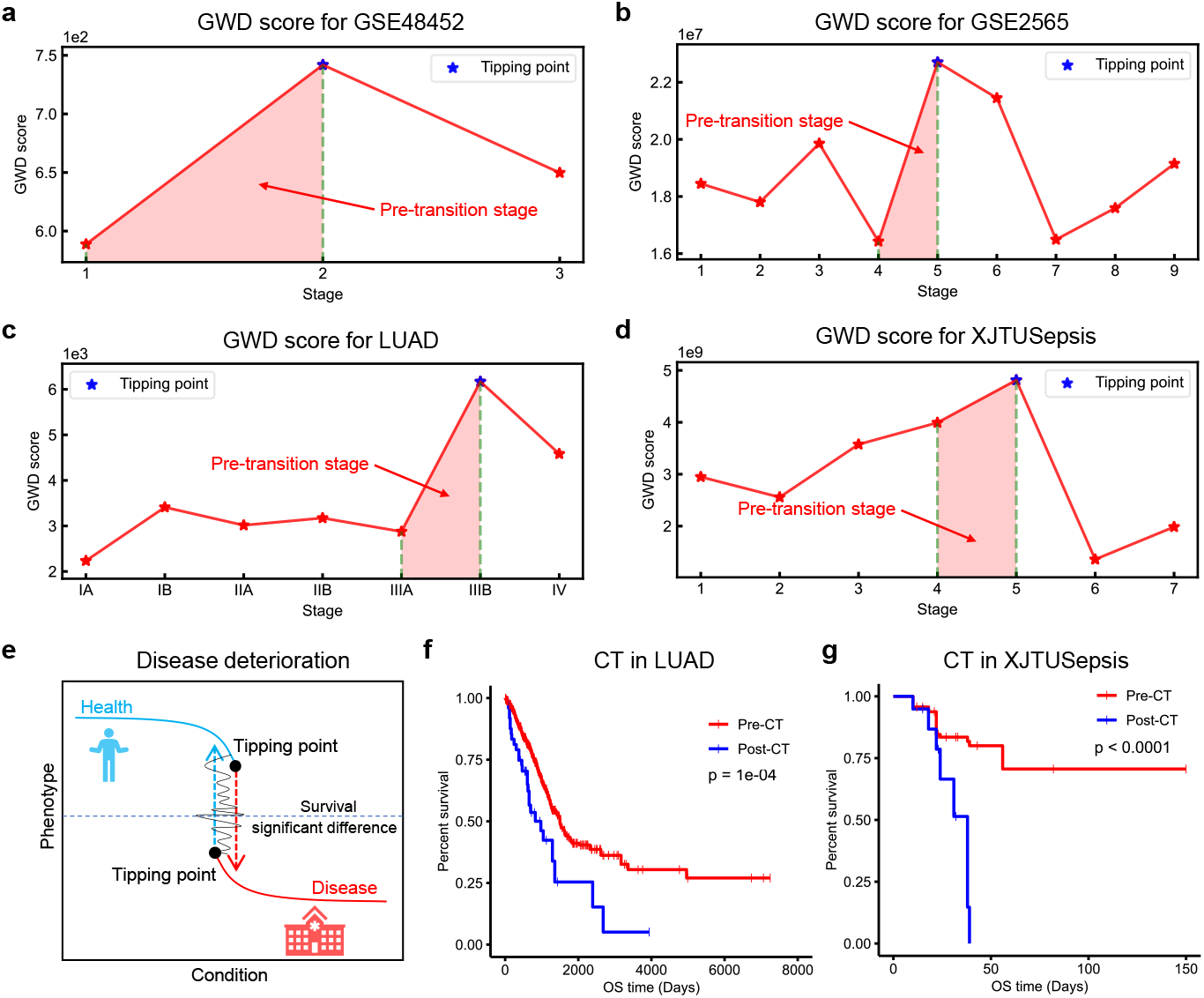
GGOT detects critical transitions in diseases with varying progression rates. The curve of GWD scores in **a**, GSE48452, **b**, GSE2565, **c**, LUAD, and **d**, XJTUSepsis to detect critical transitions. GWD scores directly reflect critical changes in gene networks. The sharp increase in GWD scores suggests the upcoming tipping point and the critical transition may occur. The stage closest to the tipping point is called the pre-transition stage. Patients in the pre-transition stages are very susceptible to deterioration. **e** The significant difference in patient survival time before and after critical transitions (CT). Survival analysis for critical transitions in **f**, LUAD and **g**, XJTUSepsis is performed to demonstrate the validity of these critical transitions. The survival time of patients before critical transitions is significantly longer than the time after critical transitions (p ≤ 0.0001).

Non-alcoholic fatty liver disease (NAFLD) is a chronic progressive non-critical disease that usually begins with simple fatty liver (accumulation of fat in the liver) and may progress to non-alcoholic steatohepatitis (NASH, hepatocyte destruction, inflammation generation). GSE48452 consists of four stages, i.e., control, healthy obese, steatosis, and NASH, to analyze NAFLD. GGOT detects the tipping point of GSE48452 as the stage 2 “steatosis” (Fig. 3a). Before stage 2, Liver damage is reversible, and normal liver function can be restored if measures are taken. But after stage 2, the sudden decrease in lipid correlations between blood and liver indicates the occurrence of an irreversible change [47, 48].

GSE2565 is employed to investigate the mechanism of lung injury, comprising ten stages determined by the time of exposure to phosgene, i.e., control, 0h, 0.5h, 1h, 4h, 8h, 12h, 24h, 48h, 72h. Lung injury is an acute progressive critical disease. GWD scores of GSE2565 demonstrate a significant increase from stage 4 to stage 5, confirming that critical transition of acute lung injury occurs at stage 5 (Fig. 3b). The disease undergoes sudden deterioration after stage 5, with 50% − 60% of the mice succumbing in stage 6, indicating the presence of irreversible changes beyond the tipping point. This phenomenon is consistent with the results of detection [49].

Lung adenocarcinoma (LUAD) and colon adenocarcinoma (COAD) are chronic progressive non-critical diseases, with corresponding pathologic stages. The tipping point of LUAD is detected in stage IIIB, signaling the critical transition into the cancer metastasis stage (stage IV) of lung adenocarcinoma (Fig. 3c). The tumor cells invade distant tissues of other organs at stage IV, usually called advanced or metastatic cancer. After stage IIIB, the disease state deteriorates in patients, and the survival rate of patients is reduced. Critical transitions can significantly differentiate the disease states of patients (Fig. 3e). We evaluate a survival analysis of obtained patients with LUAD and compare survival curves for samples before and after the tipping point. The survival time of patients before stage IIIB is significantly longer than patients after stage IIIB (Fig. 3f). The detection signifies that GGOT serves as an early warning for cancer metastasis, and the result of COAD is similar (Supplementary Fig. 4a).

Sepsis is an acute progressive critical disease, and the previous methods can not capture the state of its progression. We collect gene expression data on sepsis from the First Affiliated Hospital of Xi’an Jiaotong University, named XJTUSepsis, and group the patients based on the SOFA scores [50] into eight stages. GGOT effectively identifies the critical transition of sepsis at stage 5 (Fig. 3d). The SOFA score interval corresponding to stage 5 is [12, 14] (Supplementary Table 4), after which sepsis severely worsens, accompanied by the impact or failure of multiple organs. The patients may rapidly transition from a stable, critically diseased state to an extremely dangerous state. The corresponding survival analysis also demonstrates the accuracy of the sepsis tipping point (Fig. 3g). GSE154918 is another data about sepsis and consists of four stages, i.e., healthy control, uncomplicated infection, sepsis, and septic shock. Overall, the GWD score curve of GSE154918 is shown in Supplementary Fig. 5a. GGOT recognizes the transition signals at the early stage of the disease state transition, which can help doctors intervene early to achieve the role of delaying disease progressions or reversing disease states.

### 2.4 GGOT identifies trigger molecules to uncover the disease progression mechanisms

GGOT allows us to isolate the role of the single gene, named the LWD score, by decomposing the GWD score. The LWD scores for different diseases are illustrated in Figs. 4a-d, Supplementary Figs. 4b and 5b. It is observed that the LWD scores of specific molecules increase dramatically near tipping points, while the LWD scores of the others remain low throughout whole disease progressions. The landscapes of LWD scores in chronic diseases exhibit slower changes compared to those in acute diseases. The molecules with high LWD scores at the tipping point stage are the graph components that induce global structural changes, called the trigger molecules. These trigger molecules are more likely to play a key role in disease progression.

**Fig. 4:**
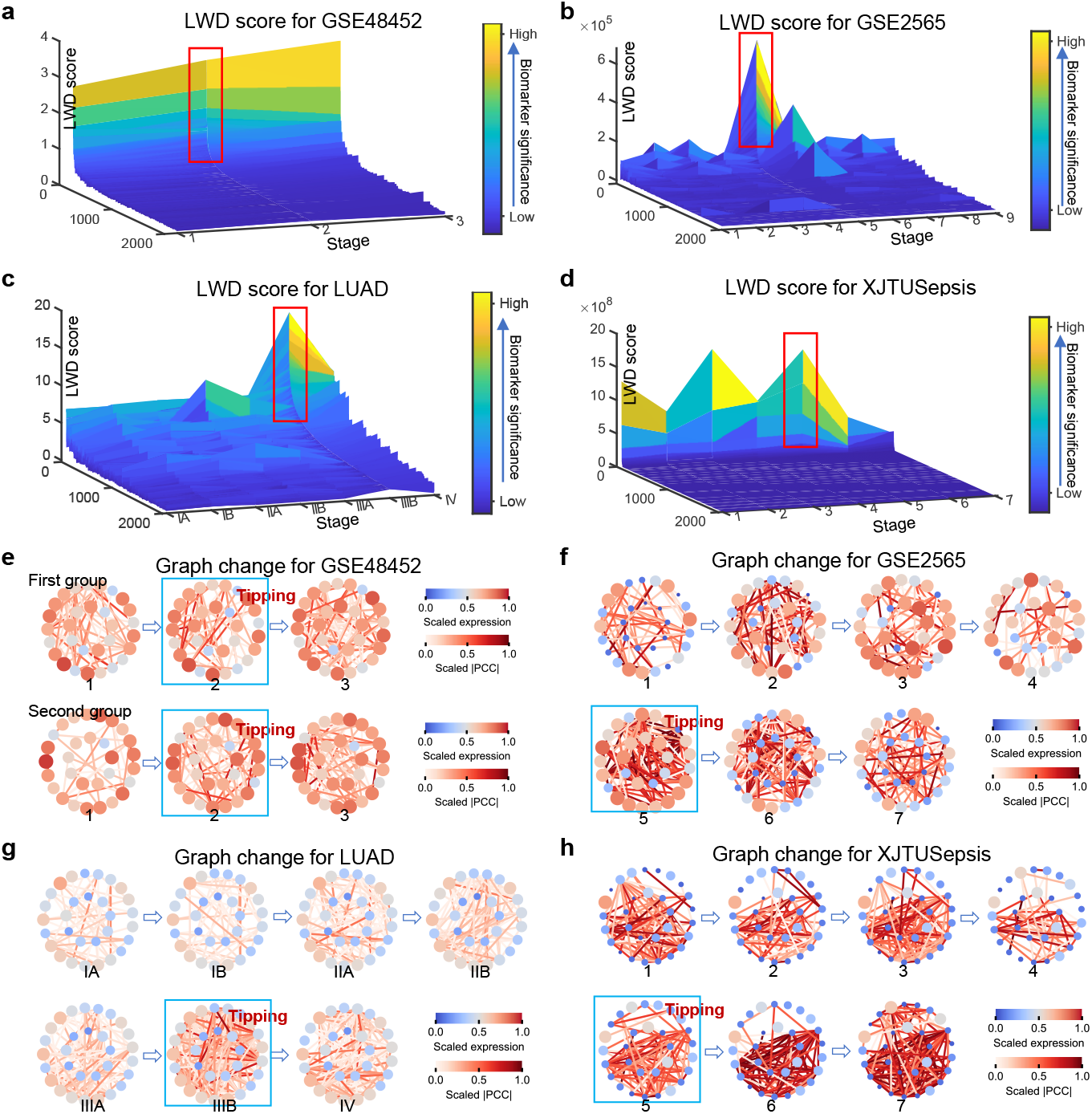
GGOT identifies trigger molecules during disease progression, revealing mechanisms of interaction relationship changes in trigger molecules. The landscape of LWD scores in **a**, GSE48452, **b**, GSE2565, **c**, LUAD, and **d**, XJTUSepsis to identify trigger molecules. LWD scores increase dramatically approaching the tipping point. The changes in LWD scores are mainly focused on certain molecules, called trigger molecules. The trigger molecules regulate network changes significantly and may play a greater role in disease progression. These molecules can be identified using LWD scores proposed by GGOT. We mark the trigger molecules near the critical point with red boxes. The interaction relationship changes of 30 trigger molecules in four disease datasets, **e**, GSE48452, **f**, GSE2565, **g**, LUAD, and **h**, XJTUSepsis. The size and color of nodes correspond to the respective scaled gene expression level, and the color of edges indicates the scaled Pearson Correlation Coefficient (PCC). As the sample approaches the tipping point, the associations between the trigger molecules gradually increase. The blue boxes are used to emphasize the interactions at tipping points.

We identify top 200 trigger molecules for different types of diseases in Supplementary Table 1. In addition, we show changes in the interactions of trigger molecules during disease progression (Figs. 4e-h, Supplementary Figs. 4c and 5c). The interactions between trigger molecules increase as they approach tipping points, reflecting strong correlations between trigger molecules. To further validate the rationality of the selected trigger molecules, we perform functional and survival analysis for trigger molecules of each dataset (Fig. 5). Functional analysis can substantiate molecular feasibility and elucidate the underlying mechanisms of diseases. Concurrently, survival analysis can provide insights into the prognostic implications of these molecular findings.

**Fig. 5:**
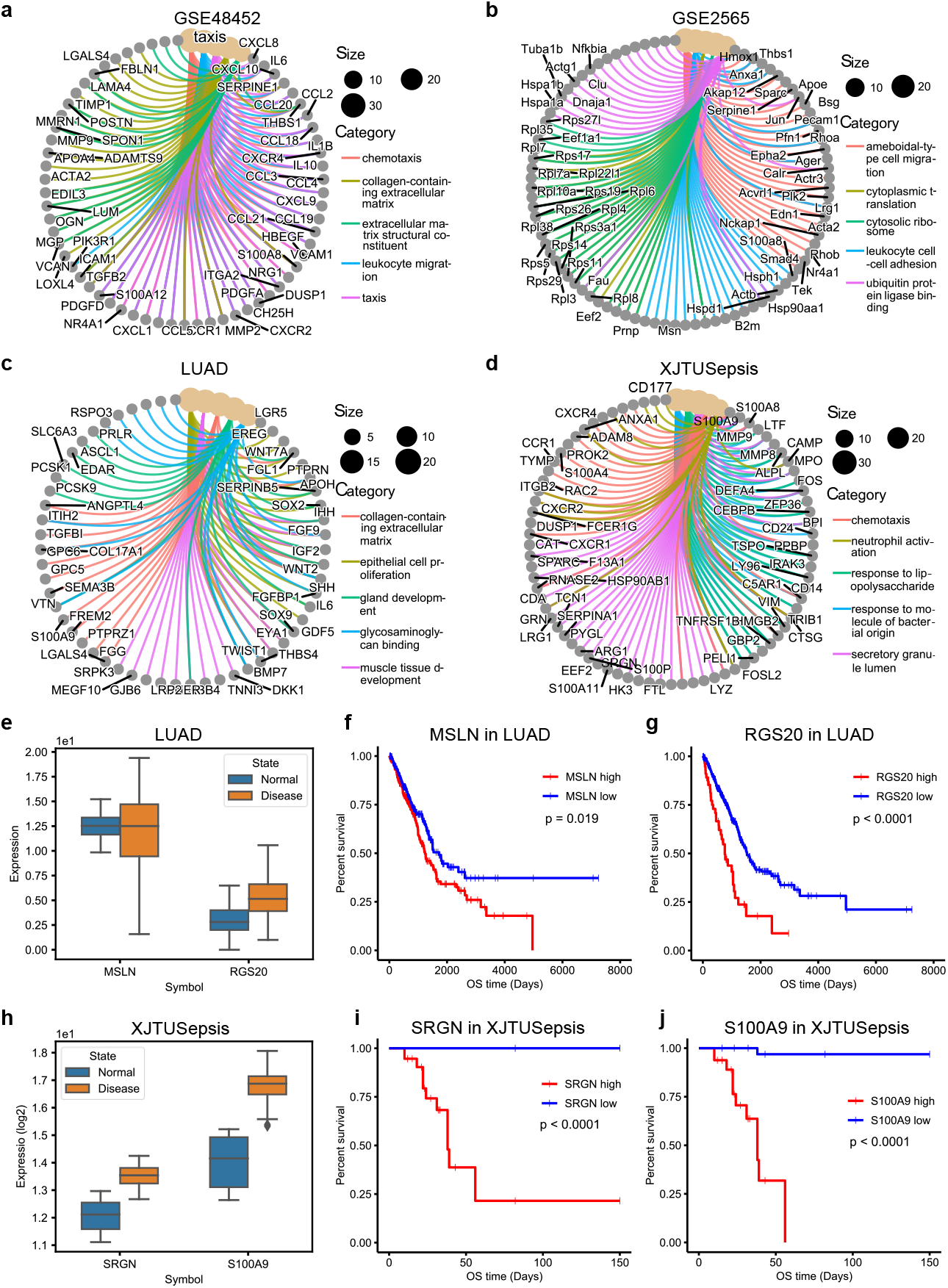
GGOT elucidates key pathways, biological processes, and prognosis in disease progressions through functional and survival analysis of trigger molecules. We perform GO and KEGG functional analysis for **a**, GSE48452, **b**, GSE2565, **c**, LUAD, and **d**, XJTUSepsis. Different colored edges indicate different functional relationships or biological processes, circles represent different genes, and node size indicates enrichment abundance, showing the association of genes with biological processes. Box line plots of **e**, LUAD and **h**, XJTUSepsis demonstrate the changes in molecular expression. The blue and yellow boxes denote normal and disease states. The survival analysis of **f**, MSLN, **g**, RGS20, **i**, SRGN, and **j**, S100A9 is performed to assess significance and understand prognosis.

GSE48452 reveals that non-alcoholic fatty liver is through biological processes of chemotaxis, leukocyte migration, etc (Fig. 5a). The molecular function of the extracellular matrix structural constituent and the cellular components of the collagen-containing extracellular matrix regulate the development of nonalcoholic steatohepatitis. The major genes constituting these processes are LAMA4, THBS1, ICAM1, the CXCL family, the CCL family, and so on, emphasizing their integral involvement in the disease’s pathogenesis.

GSE2565 suggests that lung injury is related to ameboidal-type cell migration, cytoplasmic translation, etc (Fig. 5b). Specifically, cellular components of the cytosolic ribosome and the molecular function of ubiquitin protein ligase binding influence the development of phosgene lung injury. Key genes implicated in this process, include Anxa1, Calr, Hmox1, Jun, members of the Rps family, and so on.

LUAD indicates that lung adenocarcinoma is through biological processes such as collagen-containing extracellular matrix, epithelial cell proliferation, gland development, muscle tissue development, etc. The molecular function of glycosaminoglycan binding regulates disease trajectory in lung adenocarcinoma. The major gene clusters are MSLN, RGS20, LAMA4, CXCL family, CCL family, etc (Fig. 5c). Notably, we purposely chose MLSN and RGS20 as biomarkers to validate prognostic effects (Fig. 5e). MSLN is a cell surface antigen associated with tumor invasion and is highly expressed in a variety of cancers, acting as a cell adhesion protein. MSLN plays an important role in lung carcinogenesis and epithelial-mesenchymal transition [51]. Targeting MSLN is a potential CAR-T target for many common solid tumors [52]. We find high expression of MSLN suggests a shortened survival time of patients (Fig. 5f). It has been shown that RGS20 enhances cell cohesion response, migration, invasion, and adhesion of cancer cells [53], has corresponding oncogenic potential, and is an important factor in the survival of malignant adenomas of the lung [54]. Survival analysis showed significantly longer survival time in patients with low RGS20 expression (Fig. 5g).

XJTUSepsis demonstrates the disease progression in sepsis patients is influenced by regulating biological processes such as chemotaxis, neutrophil activation, response to lipopolysaccharide, response to molecules of bacterial origin, and secretory. The main genes that activate these terms include SRGN, S100A9, MMP9, CXCR4, CD177, and others (Fig. 5d). SRGN and S100A9 have significant differences in distribution under different states (Fig. 5h). The mortality rate of patients with high expression of SRGN and S100A9 is significantly higher than that of the low expression group (Figs. 5i and 5j). The secretion of SRGN is significantly increased in LPS-activated immune cells [55], and SRGN is the only intracellular protein identified to load sulfated glycosaminoglycans. SRGN acts as a core protein covalently bound to different types of glycosaminoglycans and regulates the immune status of the body during different periods of disease development [56, 57]. S100A9 in sepsis is a calcium and zinc-binding protein, playing an important role in the regulation of inflammatory processes and immune responses [58]. Ding and Zhao et, al. demonstrate that targeted inhibition of S100A9 can reduce inflammatory responses and lung injury in sepsis [59, 60].

The results of COAD and GSE154918 are illustrated in Supplementary Figs. 4d and 5d. We show more detailed analysis results in Supplementary Figs. 6 and 7. The results show GGOT can identify critical transition trigger molecules as prognostic biomarkers and clarify the dynamic regulation control of trigger molecules, even under the noise influence of complex clinical therapeutic interventions.

### 2.5 GGOT predicts sample stage distributions, determining potential stages of unknown samples

The Gaussian graphical model of each disease stage constructed by GGOT contributes to a better understanding of disease gene interaction based on real biomolecular association networks. This is crucial for predicting whether a single sample reaches tipping points. To make predictions on unknown patients, we construct the disease stages distribution across the whole progression using Eq. (12). The stage distribution of patients provides the most direct determination of the stage at which the patient may be located. Considering the limited sample size, we use *leave-one-out* cross-validation to calculate the accuracy rate of GGOT in predicting whether the patient reaches the tipping point. It is worth noting that we are not predicting the stage that the sample is at, but predicting whether the state of the sample crosses tipping points. We show the prediction results in Figs. 6a and 6b. The numerical results are available in Supplementary Table 2. In all datasets except COAD and GSE154918, the sample prediction accuracy surpasses 0.85, and the F1 score exceeds 0.9, demonstrating the advance of the proposed GGOT.

**Fig. 6:**
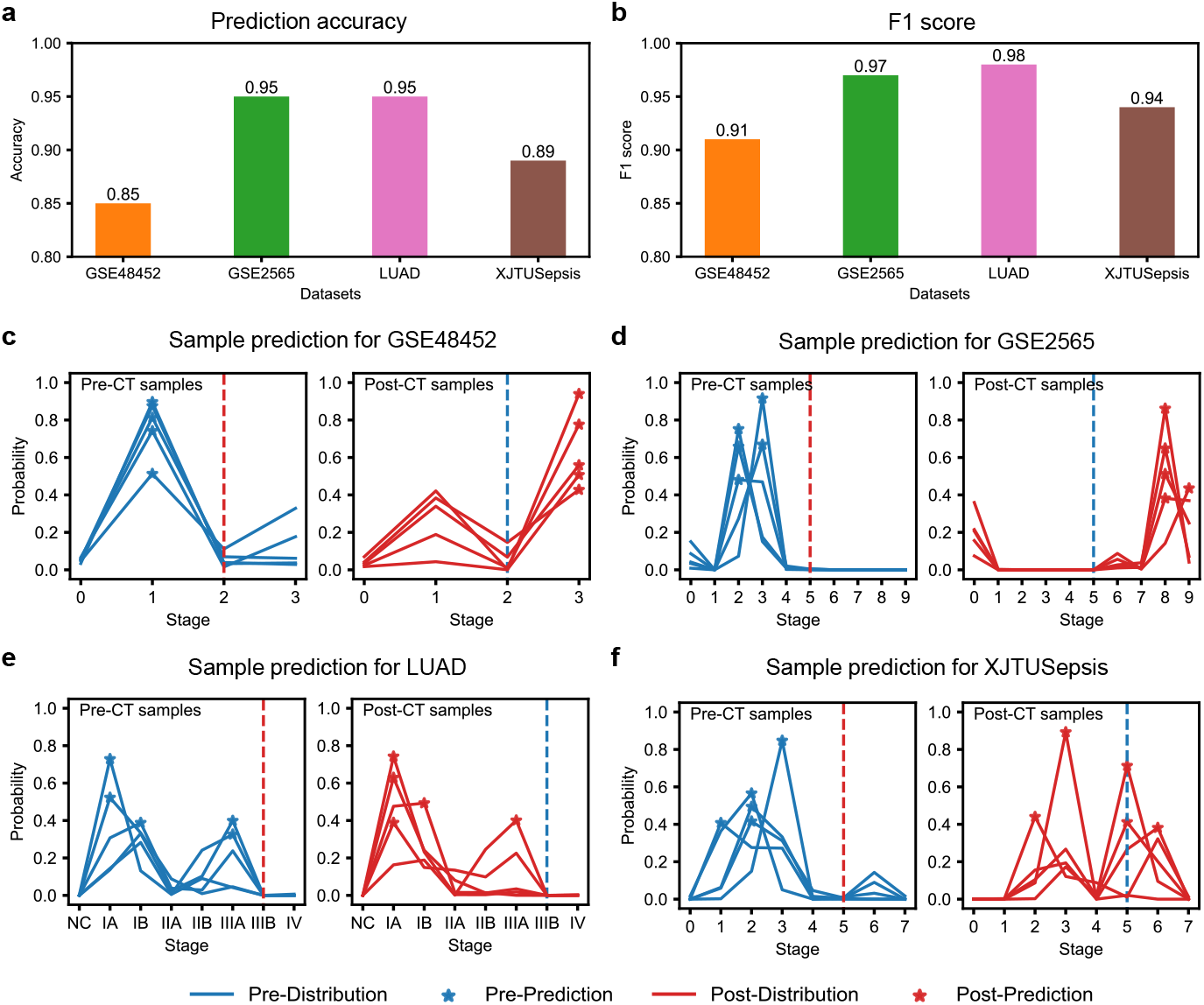
GGOT predicts whether an unknown sample reaches the tipping point. **a** The unknown sample prediction accuracies of diseases with various progression rates. **b** The F1 score of different disease prediction results. The results are evaluated from an unbalanced perspective. We further show the predictions of disease stage distribution for a single sample in **c**, GSE48452, **d**, GSE2565, **e**, LUAD, and **f**, XJTUSepsis. We denote the tipping point boundary by dashed lines. The left half is the prediction of samples before critical transitions using blue curves and the right half is the predictions after critical transitions using red curves. Different curves indicate the potential distribution of different samples, with stars representing prediction results. GGOT effectively recognizes unknown samples that are approaching the tipping point. The stage distribution of the sample is directly related to the corresponding gene expression.

We further illustrate the corresponding stage distributions of samples before and after critical transitions to analyze patient states. (Figs. 6c-f, Supplementary Figs. 4e and 5e). It is found that the prediction outcomes for non-critical disease samples are better than those for critical disease, and samples after the tipping point are prone to be mistakenly predicted as not having reached the critical state, particularly in LUAD. Critical diseases always exhibit greater complexity and the corresponding gene networks are more similar among different stages, which results in erroneous judgments by GGOT. However, on the whole, GGOT demonstrates the ability for individualized diagnosis, capable of determining if patients are experiencing irreversible transitions, aiding in personalized diagnosis and treatment.

### 2.6 GGOT visualizes stage transport processes, assessing disease progression directions

GGOT offers a novel perspective for measuring the state of disease progression, capable of accurately reflecting the phenomenon of disease critical transitions. The transport map *T* from normal to abnormal stage learned by GGOT allows us to describe the transport process of the disease progression by using Gaussian graphical distribution. The transport process of the disease is indispensable for understanding the disease progression trajectory and irreversible changes. Here, we utilize principal component analysis (PCA) [43] to demonstrate the main transformation processes in the stage distribution of different diseases (Fig. 7). The change in disease distribution is directly correlated with the GWD score. The distribution is more concentrated away from tipping points, but gradually disperses as it approaches the tipping point with the increasing GWD scores. It is demonstrated the distribution transport process of GSE48452 in Fig. 7a, and the stage state changes dramatically near tipping points. The transport processes of GSE2565 show that the distribution of stage 5 is the farthest from the normal stage (Fig. 7b), revealing that the system is on the verge of collapse. The state of disease becomes worse at the next stage 6. The state of LUAD is relatively stable before the critical transition, and difficult to diagnose. After crossing the critical transition, it deteriorates significantly with distribution changing rapidly (Fig. 7c). Sepsis progresses rapidly, making it difficult to capture the changes in stages, while GGOT can accurately describe the distribution changes in sepsis and detect that stage 5 is the tipping point (Fig. 7d). We also display the results of COAD and GSE154918 in Supplementary Figs. 4f and 5f. The more detailed results across the whole stages are shown in Supplementary Figs. 8 and 9. Meanwhile, we show distributions of trigger molecules in different stages to illustrate the regulatory changes of individual molecules in disease progression (Supplementary Figs. 10 and 11). Combining the transport processes of stage and distribution changes of trigger molecules, we find that the magnitude of state transitions in acute progressive disease is higher than the magnitude in chronic progressive diseases, which is consistent with disease progression rates. Different diseases exhibit diverse patterns of transitions during progression processes, yet GGOT can capture the alterations and detect tipping points in diseases effectively.

**Fig. 7:**
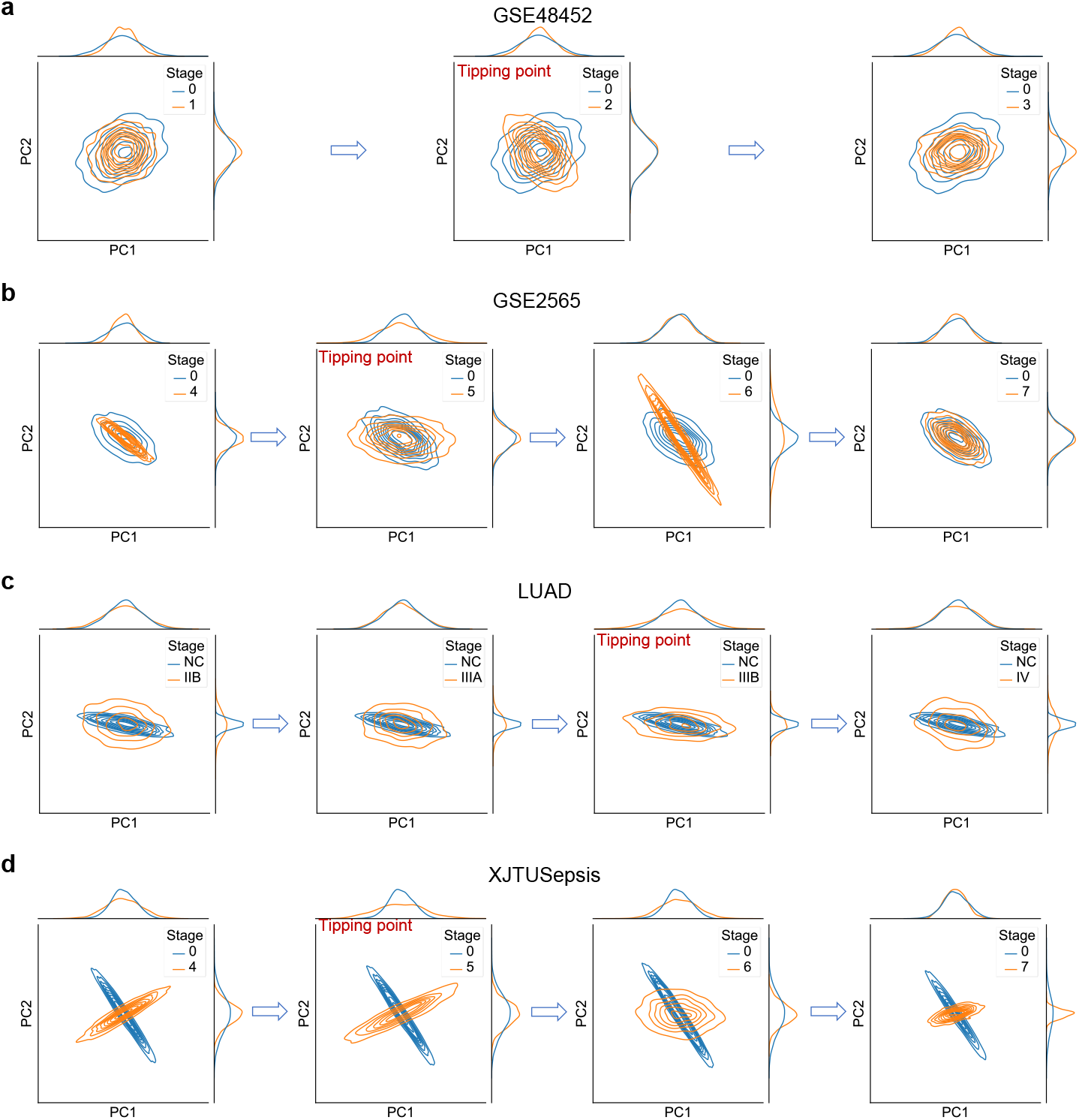
GGOT visualizes stage transport processes across disease progressions. The visualizations of the optimal transport map from normal to different abnormal stages using PCA embeddings, **a**, GSE48452, **b**, GSE2565, **c**, LUAD, and **d**, XJTUSepsis. The blue curves denote the normal stage, and the orange curves denote the abnormal stages. We focus on showcasing the changes near the tipping point. The Gaussian distribution changes little when away from tipping points, but the distribution changes dramatically near tipping points. These phenomena indicate the instability of the critical state and the suddenness of the deterioration.

## 3 Discussion

In this work, we propose Gaussian Graphical Optimal Transport (GGOT), a novel framework to model disease progression states from unpaired and unbalanced patients. By adequately modeling the nature of the problem through the lens of Gaussian graphical optimal transport, GGOT determines when the disease state reaches the tipping point, identifies the trigger molecules in critical transition, predicts the sample stage distribution, and subsequently assists in a better understanding of the distribution transport process and disease functioning mechanisms. GGOT builds on the recent successes of slowing down applications [9] and dynamical network biomarkers [11] in critical transitions, by introducing an analytic Gaussian graphical optimal transport map that can measure the relative distance of diseases’ stages as the early warning signal. Previous methods rely on statistical characteristics, and can only detect tipping points without describing the transport process of disease progression. They can not assess the importance of the location of genes in the graph structure, resulting in reduced performance in acute diseases. Besides, the limitations of data sparse and imbalance are still not resolved. Instead, we model disease progression by using optimal transport (OT) in terms of changing gene network dynamics. To the best of our knowledge, this is the first OT-based application for disease tipping point detection. GGOT combines data information and knowledge of biomolecular association networks to shape the true progression of the disease and understand the distribution of gene expression data for the disease at different stages. Distribution-based modeling makes GGOT more robust and stable, as evidenced by the strong performance of GGOT without the need for parameter tuning on various problems and different noisy datasets.

GGOT can detect critical transitions of different types of diseases without making strong assumptions about disease progressions. Unlike some approaches using differential equations for simulations, GGOT is more concerned with the information which is from the data itself. GGOT embeds protein interaction networks based on the data to enhance information characterization and remove noise. GGOT can capture the gene graph changes with progression by computing the GWD score from regulatory networks of data. The GWD score as the early warning signal is more sensitive to structural components in the graph that cause global changes, which makes it a better measure of key differences. We confirm this advantage through experiments on diseases with different rates of progression (Fig. 3). In particular, GGOT performs well in high-noise acute sepsis, demonstrating its effectiveness in acute progressive diseases.

GGOT identifies trigger molecules based on the LWD score, which quantifies the contribution of each gene to network changes. In our case, we depict the landscape of the LWD score across disease stages, observing that certain genes, identified as trigger molecules, exhibit a sharp increase in LWD scores near the tipping point. Conversely, the LWD scores of other genes consistently remain low, aligning with established findings regarding regulatory molecules (Figs. 4a-d). Besides, the correlation between trigger molecules grows when closer to the tipping point (Figs. 4e-h). The results of functional analysis and survival analysis further validate trigger molecule significance, whose enriched gene pathways reveal key factors in the progression of the corresponding diseases, while the survival analysis of the genes helps to discover biomarkers and predict the survival of patients (Fig. 5). Thus, combining GGOT with gene engineering treatments can realize precision medicine and improve patient survival.

Determining whether a single sample reaches the tipping point is significant to personal healthcare. We further analyze the distributions of different stages for a single unknown sample, which allows us to predict which stage the sample is most likely to be in and to determine the severity of the patient’s illness. In Fig. 6, we test GGOT’s ability to predict whether the sample reaches tipping points, and the results show that GGOT is highly accurate. We consider these results to suggest that the sample genes can help early detection of whether the patient is undergoing critical transitions in diseases.

We also describe the disease progression process by analyzing the optimal transport map (Fig. 7). The distribution shape changes represent the gene interaction networks changing. The most significant transition in stage distribution occurs when the disease state progressively approaches tipping points, indicating the greatest difference in tipping point relative to the normal stage. The system is at the limit of change and will undergo a critical transition. We also observe that the distribution changes are associated with the rate of disease progression. Acute progressive diseases are typically characterized by more rapid changes in distribution. In short, GGOT can assess the rate of disease progression and deterioration through changes in distribution, offering insights into disease dynamics.

We further validate the efficacy and robustness of GGOT in acutely progressive disease. Sepsis is an acute progressive critical disease with a high mortality rate. We purposely collect blood samples from sepsis patients at the First Affiliated Hospital of Xi’an Jiaotong University to obtain high-resolution time-series data (XJTUSepsis). GGOT is effective in recognizing the progression of acute sepsis and obtains similar results in XJTUSepsis and GSE154918 (Fig. 3d and Supplementary Fig. 5), suggesting that GGOT can accurately identify critical transitions in a rapidly changing process and is robust to data acquired by different technologies. There are high noises in the data due to patient heterogeneity and clinical interventions, but the detection of GGOT remains stable and valid. The above characteristics of GGOT add clinical credibility and offer the possibility of improving the treatment of pre-diseased patients. Disease progression is a complex and nonlinear physiologic process. GGOT provides a generalized framework for federating data and prior knowledge, and we can incorporate any relevant biomolecular association networks in addition to PPI networks to improve the real-world significance of the model. In a word, the use of GGOT to detect tipping points in disease provides a new way for future work, including its use to improve our understanding of disease progression, analyze molecular regulatory mechanisms, and diagnose diseases with genes. GGOT makes great contributions to the pathological analysis of diseases and individualized precision healthcare. Although GGOT is effective in detecting tipping points of disease progression, we have not adequately modeled the continuous process of disease change but rather dealt with it discretely, which is an area that can be improved in the future.

## 4 Methods

The details of the theoretical background and related work can be found in Supplementary Section A.

### 4.1 Preliminaries

#### 4.1.1 Gaussian graphical model

Gaussian graphical models [37] are probabilistic graphical models, which can represent the conditional dependencies between variables through Gaussian distributions [61], e.g., the genetic association networks [38] for modeling relations or dependencies of genetic factors. Indeed, the Gaussian graphical model is constituted by a graph structure coupled with a Gaussian distribution.

The random vector *X* ∈ ℝ^*d*^ follows the *multivariate Gaussian distribution* 𝒩(*μ*, Σ) with mean vector *μ* ∈ ℝ^*d*^ and covariance matrix Σ ∈ ℝ^*d*×*d*^, where Σ is a symmetric positive semi-definite matrix. The corresponding density function is

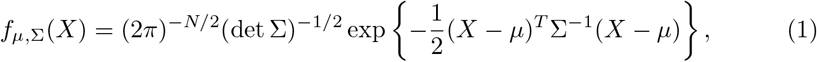

where *X* ∈ ℝ^*d*^, and *d* denotes the number of variables in *X*. In addition, we denote a graph by 𝒢 = (*V, E*) with *V* representing *d* variables in *X* and *E* denoting the set of edges among these variables. Each edge indicates the dependency between two variables.

The Gaussian graphical model is based on the multivariate Gaussian distribution [61], but utilizes the graph to depict the dependency among variables in the multivariate Gaussian distribution of Eq. (1). A random vector *X* ∈ ℝ^*d*^ is said to satisfy the *Gaussian graphical model* with graph 𝒢, if *X* has a multivariate Gaussian distribution 𝒩(*μ*, Σ) with Σ^−1^(*i, j*) = 0 for all (*i, j*) ∉ *E* [37]. Obviously, graph 𝒢 describes the sparsity pattern of the precision matrix Σ^−1^. It shows that conditional independence relations in the Gaussian graphical model correspond to the missing edges in 𝒢.

#### 4.1.2 Optimal transport

Optimal transport is a theory about distribution transport, which plays a key role in transcriptomic data analysis [41, 42]. It introduces a mathematically well-characterized distance metric, i.e., Wasserstein distance, between distributions as well as provides a geometry-based approach to realize couplings between two probability distributions [36]. This distance measure can be used to analyze the stability of the system state, which is instructive for critical transition warnings. Let *ν*_1_ and *ν*_2_ be two measures in ℝ ^*d*^. The Wasserstein distance between *ν*_1_ and *ν*_2_ [36] based on optimal transport is defined as

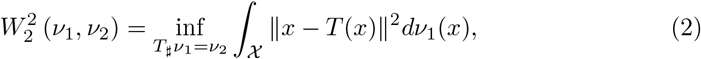

where *T* is the optimal transport map corresponding to the smallest “cost” on a metric space *𝒳*, and 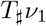 denotes the push-forward operation from *ν*_1_ to *ν*_2_. This formulation is non-convex and challenging to solve. However, when *ν*_1_ and *ν*_2_ are Gaussian distributions with zeros mean and Σ_1_ and Σ_2_ as covariances, i.e.,

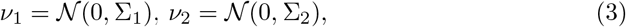

the 2-Wasserstein distance can be written explicitly in terms of covariance matrices as

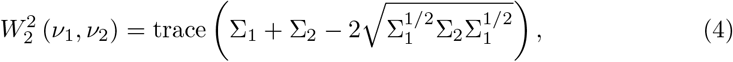

and the optimal transportation map is 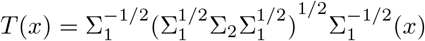.

The Wasserstein distance captures the distributional changes of the measures, and it can model the changes in structural information of the underlying graph represented by the covariance matrix. It is more sensitive to graphical differences that lead to global changes, rather than the differences that have little influence on the graph changes [62]. It is known that the gene regulatory network undergoes global changes before irreversible deterioration of the disease occurs [31]. As a result, for the gene regulatory network of a disease, this capability enables the discovery of critical transitions during the progression of the disease. The Wasserstein distance can effectively identify differences in network components by comparing the disease network to a normal network, which can be used to detect the critical transitions and identify the trigger molecules that contribute to the critical abrupt mutation causing the disease. Moreover, the optimal transport map *T* enables the movement of the disease stage from one gene graph to another, which is important for predicting the gene graph of the next disease stage. We can directly describe the complex progression of diseases by optimal transport map.

### 4.2 The Gaussian Graphical Optimal Transport Model

Recent high-throughput technologies provide a more in-depth understanding of disease progression. However, these data are often unbalanced in sample size across disease stages and lack temporal resolution and alignment. The disease samples are noisy, and can not necessarily provide all the information about disease progression in individual patients. In the following, we describe our approach, *Gaussian Graphical Optimal Transport* (GGOT) model, that detects disease critical transitions by measuring the difference between normal and abnormal stages using optimal transport, and each stage of the disease is modeled as Gaussian graphical distribution.

#### 4.2.1 Constructing gene graph associated with PPI networks

The disease progression is regulated by gene interaction networks, with different stages of the disease being determined by different gene interaction networks. So we characterize the gene interaction network as a graph to depict the disease stage. The variables *V, d*, and *E* respectively denote the set of genes, the number of genes, and the gene interactions in our problems. The edge (*i, j*) ∈ *E* implies that there is a regulatory relationship between genes *i* and *j*. The graphs are described using the Gaussian graphical model. Assuming that the gene network at a particular stage of the disease follows a Gaussian distribution 𝒩(*μ*, Σ), we are given *n* independent and identically distributed observations *X*^(1)^, …, *X*^(*n*)^ from 𝒩(*μ*, Σ). The corresponding log-likelihood can be written as

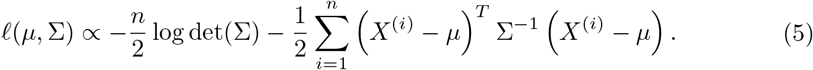

The unbiased estimate of (*μ*, Σ) is derived by 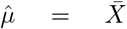 and 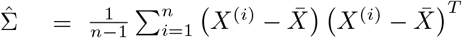, where 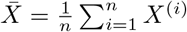. The *μ* reflects the mean gene expression levels at the current stage, and the Σ characterizes the interactions between genes and determines the structure of the graph. Specifically, The covariance matrix elements Σ(*i, j*) determine the global correlation between gene *i* and gene *j*, with Σ(*i, j*) = 0 indicating that gene *i* and *j* are globally independent. Meanwhile, the precision matrix elements Σ^−1^(*i, j*) reflect the local correlation between genes *i* and *j*, where Σ^−1^(*i, j*) = 0 signifies that genes *i* and *j* are locally independent after giving the remaining genes.

According to the dynamic changes of diseases and the validation of clinical information, we propose that the disease progresses through *N* + 1 stages, with Ω_*S*_ = {0, 1, …, *N*} denoting the disease stage index set. Given a temporal gene expression dataset 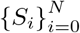 encompassing *N* + 1 stages for the disease, where *S*_*i*_ represents the set of disease samples at stage *i* containing *n*_*i*_ samples, we assume that *S*_0_ corresponds to the sample set of patients in the normal stage and the subsequent stages *S*_*i*_ for *i >* 0 are indicative of various abnormal stages. The size *n*_*i*_ of stage *S*_*i*_ varies with the index *i*, and 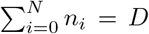. The indexes in Ω_*S*_ are ordered in increasing stages indicating the different disease progression stages. Let 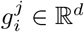 denote the sample *j* in stage *i*, 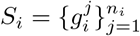, where *d* is the number of the genes. Therefore, we adopt 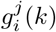 to represent the expression level of the gene *k* in the sample *j* at the stage *i*.

In our setting, the patients in different stages are not required to be aligned in identities across stages, which reduces the limitations for data. We assume the critical transition of disease is more determined by the changes in regulatory relationships between genes rather than the ones in the gene expression values. To eliminate the expression variation effect of different stages of data and better focus on the changes of edge relations of the gene network, we centralize the data according to the stages, by subtracting the averaged gene expression value for each gene in each stage. For each stage *S*_*i*_, *i* = 0, 1, …, *N*, we define the centralized sample data 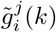 of gene *k* as

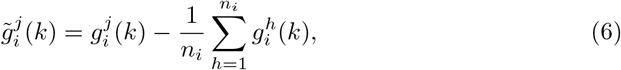

Where 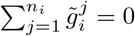. Based on this, we define a Gaussian graphical model 𝒢_*i*_ with *d* gene variables for *S*_*i*_ referred to the Section 4.1.1. It is assumed that the graph is connected, undirected, and edge-weighted. The edges characterize the interactions between genes. The corresponding Gaussian distribution of the graph 𝒢 is 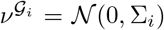 following the formulation of unbiased estimate, where 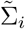 is the sample covariance as

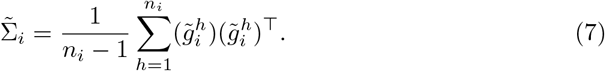

The sample covariance 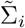 describes the gene network at the current stage of *S*_*i*_. The element 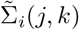 estimates the correlation between genes *j* and *k*. The magnitude of 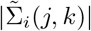 reflects the strength of the interaction between genes *i* and *j*. Furthermore, the precision matrix 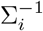 reflects conditional independence between genes. The genes *j* and *k* are locally independent if 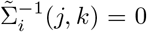, which is manifested by the absence of *j* to *k* edges in the corresponding graphs.

Moreover, the interaction of genes can be described by Protein-Protein Interaction (PPI) networks, which contribute to analyzing the phenotype of the disease. Therefore, constructing suitable protein networks using genetically related genes in complex diseases enables providing rational hypotheses for experiments. We constructed the corresponding disease PPI global network, denoted as *P*, utilizing the STRING database [35]. The elements of *P*(*i, j*) ∈ [0, 1] denote the interaction confidence levels of gene *i* and gene *j* by existing knowledge. We incorporate prior knowledge from the PPI network *P* into the covariance matrix 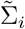 to enhance its descriptive ability for the real gene network, eliminating redundant relationships within the network as shown in Fig. 1c. The covariance matrix Σ_*i*_ for the Gaussian distribution of the disease, based on the real biological significance, is defined as

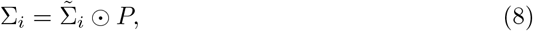

where ⊙ is the Hadamard product of the matrix. The Σ_*i*_ provides a better approximation to the true distribution of the corresponding disease stage by taking into account both data information and biological prior knowledge.

#### 4.2.2 Detecting critical transitions via optimal transport

To find an effective strategy for analyzing differences across various disease stages, we build the optimal transport maps based on normal (*i* = 0) and abnormal (*i* ≠ 0) stages. For each stage *i* ∈ Ω_*S*_, we can obtain a Gaussian graphical model 𝒢_*i*_ whose distribution is 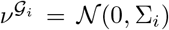. We interpret graphs as key elements that drive the probability distributions of genes in different stages. Instead of comparing patients’ gene expressions, we concentrate on the gene distributions in different stages, which are governed by the graphs. Meanwhile, optimal transport can find the minimum distance between normal and abnormal distributions. The disease state dissimilarity between normal graph 𝒢_0_ and abnormal graphs 𝒢_*i*_(*i* = 0) is measured through the Wasserstein distance (Fig. 1d). It is defined as the **Global Wasserstein Distance** (GWD) score *G*, i.e.,

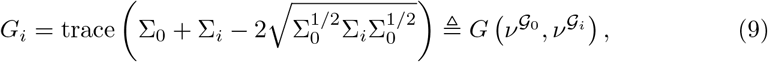

where *G*_*i*_ reflects the minimum “effort” required to recover from abnormal state *i* to normal state. The corresponding optimal transport map 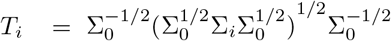 denotes the push forward of 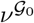 to 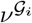. The GWD score captures the structural information of the entire graph under comparison, and it is highly sensitive to differences resulting in global changes in comparison to directly comparing covariance. This allows it to precisely analyze the most significant differences in networks across different disease stages which is important for detecting the tipping point.

The framework described above allows us to establish maps *T*_*i*_ from normal (*i* = 0) to abnormal stages (*i* ≠ 0). The corresponding *G*_*i*_ reflects differences during stage changes, which can serve as an early warning signal for disease-critical transition. As the disease state approaches the tipping point, the dissimilarity of the disease network increases, i.e., the *G*_*i*_ enlarges. The internal stability of the system deteriorates at the juncture. When far from the tipping point, the dissimilarity of the disease network diminishes. Hence, we detect the tipping point *I* ∈ Ω_*S*_ of the complex disease as

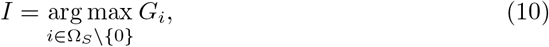

where the time point *I* reveals the time at which the disease reaches the critical transition point, during which the gene interaction network of the disease exhibits the maximum difference compared to the normal stage (Fig. 1e). The variations in graph components, such as the increased gene variance, and the increased correlation between genes, lead to global changes in the gene network. To further validate the effectiveness of our method, we performed a survival analysis, and the results can be found in Section 2.3.

#### 4.3.3 Identifying trigger molecules in transitions

Gaussian graphical model embedded with PPI network enables the gene graph 𝒢_*i*_ to describe the gene interactions of the disease. However, Wasserstein distance only reflects the whole difference of complex interactions at different disease stages. Identifying the key regulatory genes during the critical transition period of disease occurrence is more crucial. Indeed, we can decompose the Wasserstein distance into gene-based **Local Wasserstein Distances** (LWD) score *L* by considering the complex interaction of individual genes with other genes.

The formulation of Wasserstein distance in Eq. (9) is mainly carried out by the covariance, which is related to the corresponding graph structure. The LWD score of stage *i* is defined as

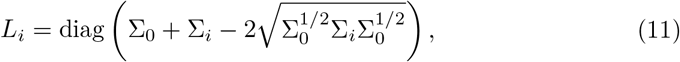

where *L*_*i*_ ∈ ℝ^*d*^, indicating the whole of the interaction of each single gene. We let 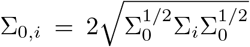, where Σ_0,*i*_ ∈ ℝ^*d*×*d*^ is a symmetric positive definite square matrix. Supposing that *σ*_0_(*j*), *σ*_*i*_(*j*), *σ*_0,*i*_(*j*) represent the *j*-th diagonal element in Σ_0_, Σ_*i*_, Σ_0,*i*_ respectively, we rewrite the LWD score of gene *j* at stage *i* as *L*_*i*_(*j*) = *σ*_0_(*j*) + *σ*_*i*_(*j*) *− σ*_0,*i*_(*j*), where *L*_*i*_(*j*) is the *j*-th element of *L*_*i*_. The GWD score *G*_*i*_ can be decomposed as the sum of *L*_*i*_(*j*), i.e., 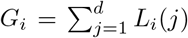, where *L*_*i*_(*j*) reflects the difference in distribution caused by the gene *j*. The previous methods only calculate changes in gene interactions, ignoring the importance of gene location in the graph structure. However, we find the score *L*_*i*_(*j*) considers the importance of gene location when computing gene interactions. We demonstrate this conclusion by controlling the strength of the interactions and changing the position of the nodes (Supplementary Fig. 1).

Considering the LWD score at tipping point *I*, the *L*_*I*_ (*j*) denotes the contribution degree of gene *j* to the critical transition phenomenon at the tipping points. The significance of gene *j* is gauged by the magnitude of |*L*_*I*_ (*j*)|, serving as a criterion to identify trigger molecules. In light of this, we can effectively identify trigger molecules by screening the top *C* molecules with larger LWD scores that cause the critical transition of the disease. The trigger molecules are used for the downstream analysis, which aids in disease diagnosis and gene therapy.

To further validate the regulatory mechanisms of the trigger molecules, we perform the gene functional analysis. Gene functional analysis is the process of categorizing genes according to gene prior knowledge, i.e., genome annotation information. The functional analysis including gene ontology and pathway enrichment was based on GO database and KEGG database. Gene Ontology [63, 64] is a functional database of the computational knowledge structure of genes, including Biological Process (BP), Molecular Function (MF), and Cellular Component (CC). GO annotation helps to understand the biological function and significance of selected expressed genes. KEGG Pathway Enrichment [65], based on biological pathways, enables the identification of the biochemical metabolic pathways and signaling pathways involved in selected expressed genes. The results are available in Section 2.4.

#### 4.2.4 Predicting sample distributions for early diagnosis

Determining whether a patient reaches the critical transition point of the disease is crucial for early intervention. In addition to predicting tipping points, our model is capable of forecasting the stage distribution of unknown samples by parameterizing Gaussian graphical distributions for *N* abnormal stages. For a sample *s* ∈ ℝ^*d*^ with unknown stage, we define the probability *p*_*i*_ of this sample being in stage *i* as

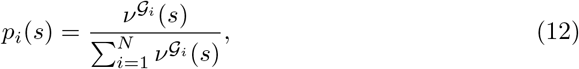

where 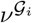 is the probability density function of the Gaussian graphical distribution of stage *i*. The probability *p*_*i*_ reflects the relative probability of the sample potential stage *i* in the Euclidean space ℝ^*d*^. Therefore, the true stage *I*_*s*_ that sample *s* belongs to is predicted as

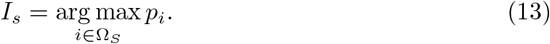

Stage *I*_*s*_ is the period where the potential stage probability for sample *s* reaches its maximum. It determines the predicted stage for the sample, providing assessments of the disease severity and whether the tipping point has been reached. Due to the limited number of samples, we employ *leave-one-out* cross-validation for single sample stage prediction to assess the effectiveness of our method. We reserve one data point from the available dataset for prediction and train the model based on the remaining data. After repeating the experiment for *D* times, we compute statistics on the accuracy of the model predictions (Section 2.5).

#### 4.2.5 Visualizing disease transport processes across progressions

GGOT utilizes optimal transport to establish the distribution transformation from normal to abnormal stages of diseases. The transformation is induced by changes in gene interactions. Because the number of genes associated with the disease is extremely large, it is challenging to describe stage change trajectories in diseases under high-dimensional gene data. The optimal transport map *T*_*i*_ describes the transport process of the Gaussian distribution from 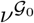 to 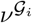. We can redefine transport maps using principal components or trigger molecules to represent the global or local distribution changes, as *T*_*i*_ is analytical. For the global transport process of disease, we employ principal component analysis to retain the top two components in gene samples. After data dimensionality reduction, the resulting covariance matrix is Σ′ ∈ ℝ^2×2^. The corresponding low dimensional optimal transport map 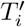 from 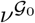 to 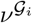 is represented by linear transformation. We can look across the global progression of disease progression via 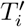. As for the trigger molecules stage changes, we consider the marginal distribution 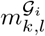 of gene pair (*k, l*). We find 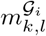 is still following the Gaussian distribution as

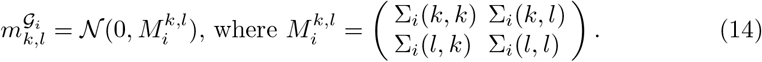

The 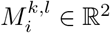 denotes the covariance of gene pair (*k, l*). The stage changes in selected trigger molecules (*k, l*) can be accurately characterized by 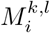. We analyze changes in the relationships between several genes through the marginal distribution, unaffected by the influence of the global genes. The results are shown in Section 2.6.

### 4.3 Datasets and preprocessing

We apply the GGOT method to six time-course or stage-course datasets, i.e., the lung adenocarcinoma (LUAD), the colon adenocarcinoma (COAD) from [TCGA](https://portal.gdc.cancer.gov/) database, the non-alcoholic fatty liver (GSE48452), the lung injury (GSE2565), the sepsis (GSE154918) from [GEO](https://www.ncbi.nlm.nih.gov/geo/) database, and our collected dataset of sepsis patients from the First Affiliated Hospital of Xi’an Jiaotong University. For all real-world datasets, we discard the probes without corresponding NCBI Entrez gene symbols. Meanwhile, for each gene symbol mapped by multiple probes, the maximum or average value is employed as the gene expressions. The procedures of selecting sample and preprocessing data are as follows. First, we screen suitable patient samples based on clinical information and disease progression status. The screened patients should conform to the progression direction of the disease. Second, we define two parameters *r* (expression rate) and *h* (high expression level) to select highly expressed genes in samples. The gene *k* is chosen if the proportion of samples with gene *k* exhibiting an expression level higher than *h* is at least *r*. This approach effectively eliminates genes with low or no expression in the samples, reducing the impact of irrelevant variables in the experiment. Then, the Protein-Protein Interaction networks for Homo sapiens and Mus musculus are downloaded from [STRING](https://string-db.org/) database [35]. We integrate this information into the largest global gene graph *P*_*h*_ for the respective species. Last, we map the genes from each dataset to the corresponding global gene graph, representing gene interactions based on prior knowledge for consequent analysis. When mapping the graph, we set the threshold *u* of confidence level. The edges with gene confidence levels greater than *u* are retained, and the genes corresponding to isolated nodes will be removed. Additionally, we conduct differential gene analysis between abnormal and normal groups to assist in selecting trigger molecules. The experiment parameters are available in Supplementary Section E.

## 5 Data availability

The details of the simulation data are available in Supplementary Section D. Raw published data for the non-alcoholic fatty liver [66], the lung injury [49], the sepsis [67] are available from the Gene Expression Omnibus ([GEO](https://www.ncbi.nlm.nih.gov/geo/)) under accession codes GSE48452, GSE2565, GSE154918, respectively. The lung adenocarcinoma and the colon adenocarcinoma datasets are from the Cancer Genome Atlas Program ([TCGA](https://portal.gdc.cancer.gov/)). Their original data are downloaded at the link https://portal.gdc.cancer.gov/projects/TCGA-LUAD and https://portal.gdc.cancer.gov/projects/TCGA-COAD. The sources of the above data are provided with this paper. The Access of XJTUSepsis about sepsis from the First Affiliated Hospital of Xi’an Jiaotong University requires authorization. The processing data sets for all tasks can be accessed by contacting.

## 6 Code availability

The GGOT method is written in Python and uses standard Python libraries, for detecting critical transitions and identifying trigger molecules to understand disease progression. The source code of our proposed GGOT is available at https://github.com/huawenbo/GGOT.

## 7 Acknowledgments

This work was supported by National Key R&D Program 2021YFA1003000, National Natural Science Foundation of China 12125104, and Shannxi University Joint Program 2021GXLH-Z-099. We thank the Department of SICU of the First Affiliated Hospital of Xi’an Jiaotong University, Department of Hepatobiliary Surgery and Liver Transplantation of the Second Affiliated Hospital of Xi’an Jiaotong University, and Key Laboratory of Surgical Critical Care and Life Support of Xi’an Jiaotong University Ministry of Education for their support and collecting data. We also acknowledge [GEO](https://www.ncbi.nlm.nih.gov/geo/) and [TCGA](https://portal.gdc.cancer.gov/) databases for providing their platforms and contributors for uploading their original datasets.

## 8 Author contributions

W.H., R.C and J.S. conceived the method. W.H. implemented the method. W.H. and J.S. generate the numerical results. W.H., R.C. generated the experimental results. W.H., R.C., J.S., C.L. interpreted the results. W.H., R.C., H.Y. and J.Z. generated the diagrams and wrote the paper. All authors reviewed the manuscript.

## 10 Additional information

The additional information is available at **Supplementary Information**.

